# An index of the quality of evidence in meta-analysis of randomized controlled trials

**DOI:** 10.1101/100081

**Authors:** Devon D. Brewer

## Abstract

The quality of evidence in meta-analysis of randomized controlled trials is the degree to which the estimated effect represents the “truth.” Current approaches to assessing the quality of evidence focus on trial design and methods. I describe a new quality of evidence index composed of four sub-indexes that measure pre-registration, independent replication, data availability, and trial design and methods, respectively. This index is systematic, objective, and quantitative. I illustrate the index with an empirical example and provide a spreadsheet for easy calculation.

> *“…when you can measure what you are speaking about, and express it in numbers, you know something about it; but when you cannot express it in numbers, your knowledge is of a meagre and unsatisfactory kind.”*
>
> — *-Lord Kelvin (William Thompson), 1824-1907*

## Introduction

Meta-analyses of randomized controlled trials (RCTs) typically involve objective, quantitative summaries of at least four aspects of the evidence: amount (numbers of trials and participants), an estimated effect, precision of the estimate, and consistency of results. These aspects can be summarized concisely and, for the latter three, graphically, allowing easy interpretation even by non-scientists. However, many meta-analyses of RCTs do not indicate in objective, quantitative terms the quality of the evidence, that is, the degree to which the estimated effect represents the “truth.” The Grading of Recommendations Assessment, Development, and Evaluation (GRADE) working group defined quality of evidence similarly: “… the extent of confidence that an estimate of effect is correct” (p. 995) (1). The quality of evidence is determined by how the evidence was produced, regardless of the results observed. It is logically distinct from the magnitude of the estimated effect, numbers of trials and participants, consistency of results across trials, and precision of the estimated effect.

Existing approaches to assessing the quality of RCTs have mostly focused on trial design and methods, such as blinding, concealment of the randomization sequence, and intention-to-treat analysis (2–4). Many researchers label the absence of one or more of these features as *bias*, even though such flaws do not necessarily shift results in a particular direction. Nonetheless, flaws in trial design and methods do reduce confidence in results, and thus decrease the quality of evidence. The GRADE approach to assessing quality of evidence also entails informal consideration of publication bias (the possibility that unreported trials may have different results than reported trials) (5) as well as properties logically separate from quality (2), including the magnitude of the estimated effect, precision of the estimate, and consistency of results (4).

There seems to be no disagreement that characteristics of trials can be assessed objectively and that certain characteristics indicate either higher or lower quality of evidence. Nonetheless, in some approaches to assessing quality of evidence, evaluations are not standardized or combined systematically and formally (2, 4). Other approaches, however, do involve quantitative measures of quality (3).

Some researchers have examined the relationship between quality scores and estimated effects in particular sets of trials, and then judged the utility of assessing quality by the magnitude and direction of these associations (6–9). These authors did not articulate why they thought quality might be related to trial results. In my opinion, such investigations cannot reveal the utility of assessing quality, as estimated effects bear no necessary relationship to quality. Trial design and methods may influence estimated effects, but in unknown and unexpected ways that vary between trials, and therefore are a source of *error.* The quality of evidence from RCTs cannot be validated empirically. Rather, quality corresponds to the degree to which the evidence was produced with procedures that reduce error (including bias) and confounding, and enhance experimental control, validity, reliability, and transparency. Quality is simply a fundamental descriptive characteristic of evidence, not a predictor of other aspects of the evidence.

I propose an index of the quality of evidence in meta-analysis of RCTs. This index includes components from prior approaches, as well as dimensions of quality that, to my knowledge, have not been summarized objectively and quantitatively in the past. While trial design and methods are critical aspects of the quality of evidence, so are pre-registration, independent replication, and data availability.

Online trial registries store study protocols for RCTs and other studies. Pre-registration occurs when researchers submit their protocols to a registry *before* enrolling participants. By declaring the intervention, comparator, participant population, and outcomes in advance, researchers demonstrate that their RCTs were conducted according to plan, and that the results they report do not stem from procedural changes, data dredging, or post-hoc hypotheses. Pre-registered RCTs also comprise a data source for direct assessment of unreported evidence.

Replication is the bedrock of science. Only through the cumulation of results from studies employing the same or similar designs and methods can reliable knowledge emerge. Confidence in results rises to the extent that researchers have worked independently from one another. Thus, independent replication is an essential component of the quality of evidence. By replication, I mean replication of similar *studies*, not necessarily replication of *results.* Consistency of results across studies is an entirely different property already well assessed with standard meta-analytic tools.

Another cardinal element of scientific practice is sharing or making public the data which underlie study results. By making their data public, researchers demonstrate, at least nominally, that they did not fabricate their results. These data enable others to reproduce the original investigators’ analyses to confirm the reported results. The public data may also allow investigation of other hypotheses not examined by the authors that are relevant to the originally reported results. In some fields, publication or sharing of data has been normative for decades or even centuries. Scientists often regard research for which the raw data are unavailable as unscientific and bar such work from the literature (10). As the Royal Society’s motto holds, “nullius in verba” (on no one’s word). And researchers’ reports are just words expressing claims.

## Description of measure

The quality index (abbreviated as Q), is a combination of four separate sub-indexes that represent quality of the evidence on each of four dimensions (pre-registration, independent replication, data availability, and trial design and methods). I detail first the calculation of each sub-index, and then describe the computation of the overall index. The quality index applies to a defined set of trials for a particular intervention, comparator, indication, and outcome that have been summarized with meta-analysis.

### Pre-registration

The proportion of trials that were pre-registered, weighted by sample size, indicates crudely the amount of evidence that was pre-registered. However, some pre-registered trials may not have been reported, even after a reasonable amount of time since they were completed. Pre-registered trials that are not reported represent known, but omitted, evidence and decrease the value of pre-registration as a signal of quality. Therefore, in calculating the pre-registration component of the index, the weighted proportion of trials that were pre-registered is reduced by the fraction of completed pre-registered trials eligible for inclusion in the review that have not been reported. I define a pre-registered but unreported trial as one that was completed more than three years before the end date of the literature search period but had not yet been reported. Three years is ample time for researchers to prepare data for analysis, conduct planned analyses, and write and publish a report. In an analysis of RCTs in the ClinicalTrials.gov registry that were completed in 2008, more than half of those that had been reported before the end of 2012 were reported within 33 months of completion (11). Formally, the pre-registration sub-index is computed as follows:

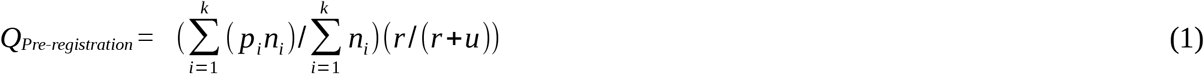

where *k* is the number of trials, *p_i_* indicates whether the *i^th^* trial was pre-registered (yes = 1, no = 0), *n_i_* is the sample size for the *i^th^* trial, *r* is the number of pre-registered trials that were reported, and *u* is the number of pre-registered but unreported trials eligible for inclusion in the review that were completed more than three years before the end of the literature search period. If all trials were pre-registered and reported, the pre-registration quality score is 1. If no trials were pre-registered or if no pre-registered trials were reported, the pre-registration quality score is 0. For assessing quality, a pre-registered trial means that the specific outcome summarized in the corresponding meta-analysis was included in the pre-registered trial protocol, and that the intervention, comparator, indication, and participants included in analysis did not change from pre-registration to reporting of results.

If the authors of a meta-analysis did not search trial registries or did not search the literature well otherwise, it is possible that another analyst computing the quality index retrospectively for that meta-analysis may find reported, pre-registered trials that were eligible for the meta-analysis yet not included in that summary. I recommend including such trials in the calculation of *r* only, even though they were not included in the authors’ original meta-analysis. Such trials indicate the original meta-analysts’ body of evidence was incomplete (and potentially biased), but they are still useful for estimating the quality signal of pre-registration for the topic at hand.

### Independent replication

The proportion of evidence produced independently, that is, by separate teams of researchers, indicates the degree of independent replication. I define a “team” here as a set of authors connected directly or indirectly by co-authors in common for trial reports included in the meta-analytic summary. In explicit terms,

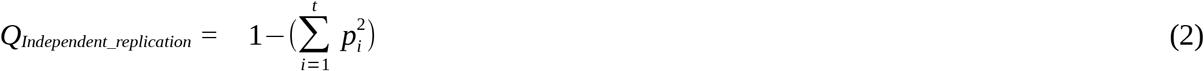

where *t* is the number of independent teams and *p_i_* is the proportion of all trial participants included in the *i^th^* team’s trials. This sub-index indicates the amount of evidence that was produced by different teams. If a single team of researchers produced all of the evidence, there has been no independent replication, and the corresponding sub-index score is 0. If, on the other extreme, the evidence is based on research produced by many independent teams with similar numbers of participants in their respective sets of trials, then there has been considerable independent replication, and the sub-index score will be a fraction relatively close to 1. This sub-index is equivalent to an index of qualitative variation that many researchers developed independently over the last 100 years (12).

Scores on this dimension, unlike other dimensions, can vary as a function of the number of trials, to the extent that each additional trial was conducted by an independent team. This is appropriate, because confidence in the accumulated evidence grows with each additional independent team’s contribution. However, there are scenarios when new evidence by a new team can reduce the degree of independent replication. Imagine several independent teams have each conducted a fairly small trial. When later a new team contributes a very large trial, the degree of independent replication likely will fall, as the large trial overshadows the other, small trials. Single large trials are never definitive and require replication just as much as small trials.

### Data availability

The data availability sub-index is the proportion of trials, weighted by sample size, for which the individual-level data have been publicly archived or published:

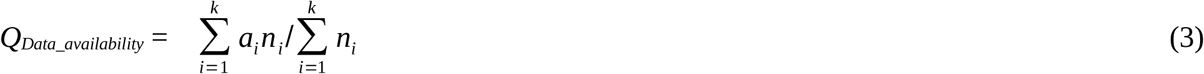

where *k* is the number of trials, *a_i_* indicates whether the data for the *i^th^* trial are publicly available (yes = 1, no = 0), and *n_i_* is the sample size for the *i^th^* trial. If authors of all trials have made their data publicly available, the sub-index score is 1. If there are no publicly available data for any of the trials, then the sub-index score is 0.

### Trial design and methods

I consider eight aspects of trial design and methods to reflect quality:

1. little differential attrition between trial arms (<= 10% absolute difference in percentages of participants assessed);
2. specified generator for random assignments to trial arms (e.g., random number table, computer, coin flips, die rolls, etc.);
3. allocation concealment (those who enrolled participants in the trial were unaware of the arm to which the next participant would be assigned);
4. units of randomization and units of analysis were the same (if individuals were randomized, then analysis focused on individuals; if groups were randomly assigned to trial arms, then analysis focused on those groups, not individuals);
5. trial participants were blinded (unaware to which arm they were assigned);
6. outcome assessors were blinded (unaware to which arm participants were assigned);
7. trial not stopped early due to the result of an interim assessment (ending the trial prematurely because of a positive or negative intervention effect at the interim assessment); and
8. intent-to-treat analysis (trial data were analyzed according to which arms participants were assigned, regardless of which interventions or how much of an intervention they actually received).

The Cochrane Collaboration (2) and GRADE investigators (4) noted most of these as factors in determining quality.

In using this sub-index, it is important to define what procedures for handling post-randomization missing data are acceptable for classifying an intent-to-treat analysis. I consider conditional mean (regression), inverse probability weighting, multiple imputation, and maximum likelihood-based imputation methods acceptable (13). If authors indicated they used intent-to-treat analysis but did not specify their imputation procedures for any missing data, I do not classify the trial as having an intent-to-treat analysis.

The trial design and methods sub-index is the mean proportion of these factors present in the included trials, weighted by sample size:

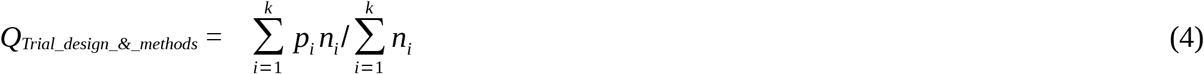

where *k* is the number of trials, *p_i_* indicates the proportion of the eight factors present in the *i^th^* trial, and *n_i_* is the sample size for the *i^th^* trial. If a trial had one or more design/methods flaws not related to the eight factors listed above, *p_i_* would decrease by 1/8 for each additional flaw. Flaws beyond the eight I noted are uncommon. Nonetheless, *p_i_* cannot take a value less than 0 no matter how many additional flaws the *i^th^* trial has. If a report does not specify whether a particular design/methods factor was present, the trial is coded as lacking that factor.

### Overall quality index

The overall quality index is a weighted mean of the four sub-indexes. I suggest that the pre-registration, data availability, and independent replication sub-indexes each be weighted 20% and the trial design and methods sub-index be weighted 40%. Although this recommendation is arbitrary, my rationale is that each component is a crucial aspect of quality and that trial design and methods deserve extra weight because they encompass several important factors. In formal terms, the overall quality index is:

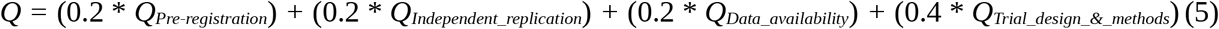

The overall index, therefore, ranges from 0 to 1, just as the sub-indexes do, with 1 representing highest quality. Index values may be simplified into a grading scheme, also arbitrary, for ease of communication. I suggest dividing the unit interval into quartiles labeled A (.75-1.0), B (.50-.79), C (.25-.49), and D (.00-.24), from high to low quality, or into duodeciles to make finer distinctions (A+, A, A-, B+, B, B-, etc.). My rationale for using the A-D scheme is to approximate the academic grading systems used in many countries, with D (or D-) often regarded as the lowest acceptable level. I consider even low quality evidence from RCTs to be better than observational evidence in nearly all circumstances.

## Example

I searched Google Scholar (scholar.google.com) in October, 2016, for a recent meta-analysis that involved use of the Cochrane risk of bias tool (2) and included fewer than 20 trials (for my ease of calculation). I chose the first I encountered which met these criteria, a meta-analysis by Machado and colleagues (14), to illustrate the quality of evidence index. Machado and colleagues (14) summarized randomized controlled trials of acetaminophen (paracetamol) as a treatment for osteoarthritis. I focused on the 8 trials (from 6 reports (15–20)) they summarized for short-term (2 weeks–3 months) pain outcomes. For my calculations, I used the sample sizes for the trials as indicated by Machado and colleagues (14). The total number of participants across trials was 2,355, and the trial reports were published between 2003 and 2014. Machado and colleagues (14) used the informal GRADE approach (4) to evaluate the quality of evidence they summarized, rating it “high quality.” A spreadsheet attached to this article shows the calculation of the quality index and the underlying information on these trials.

### Pre-registration

I searched the International Clinical Trials Registry Platform (http://apps.who.int/trialsearch/Default.aspx) in November, 2016 with the following keywords: “acetaminophen AND osteoarthritis” (searching with the keyword “paracetamol” returned the same results). I examined each of the 73 trial protocols returned from this search and kept those that met Machado and colleagues’ inclusion criteria and search dates. I found 2 pre-registered trials that satisfied these conditions, and both of them (18,20) were included in Machado and colleagues’ summary. The authors of one of these trials (18) did not indicate their trial was pre-registered (or registered at all) in their report, a fact that highlights the importance of searching trial registries. Thus, there were no eligible, pre-registered trials that had not been reported. The two pre-registered trials had samples sizes of 212 and 349, respectively, for a subtotal of 561. Dividing this by the total number of participants in the 8 trials yields a Q_Pre-registration_ value of .24.

### Independent replication

There were no overlapping authors between the 6 reports included in Machado and colleagues’ summary. However, there were two trials reported in two of the 6 reports, so at the trial level, there was author overlap. The six independent teams accounted for .02, .15, .21, .24, .30, and .09 of the participants. Therefore, the Q_Independent_replication_ score is 1–(.00 + .02 + .04 + .06 + .09 + .01) = .78.

Although it is possible to write a script to determine which sets of authors overlap, whenever the number of reports is not large (say, < 100), it may be easier to perform this task manually. One way to do this is to put the sets of authors into a spreadsheet (one set per cell) or text document (with divisions between the sets). Put the sets with the fewest authors first and then enter the rest in order of increasing number of authors. Next, search for the surnames of the authors in the first cell or division in the rest of the spreadsheet or document, respectively. If there is a match on surname and other parts of the name (and possibly other information, such as affiliation) to suggest the two names refer to the same person, then merge the two sets of authors together into the same cell or division, making clear which reports are being combined as well. Repeat this search and merge process until reaching the last set of authors (the names in which do not need to be searched). If a script is used to determine the distinct teams of authors, it is essential to inspect the results carefully for possible mistakes due to typographical errors and variations in spelling and name format in the original reports.

### Data availability

Based on the trial reports, the individual level data have not been publicly archived or published for any of the trials Machado and colleagues summarized. Thus, Q_Trial_design_&_methods_ = 0.

### Trial design and methods

I used Machado and colleagues’ determinations of whether trials had a specified generator of random assignments, allocation concealment, participant blinding, and outcome assessor blinding. I read the trial reports to assess differential attrition, units of randomization and analysis, early stopping, and intent-to-treat analysis. All trials had participant blinding, outcome assessor blinding, consistent units of randomization and analysis, and low differential attrition. No trial had been stopped early or used an intent-to-treat analysis by my definition. Authors of most trial reports did not specify the generator of random assignments or method of allocation concealment. Every trial had a majority of the design and methods quality factors present, with proportions ranging from .63 to .88. The sample size weighted mean of these proportions, Q_trial_design_&_methods_, is .68.

### Overall quality

The overall quality of evidence index for the trials summarized by Machado and colleagues is 0.48. Figure 1 shows a bar chart with the sub-index and overall index values. The evidence is of moderately high quality in terms of independent replication and trial design and methods. However, the quality of evidence is very low to low in terms of data availability and pre-registration. By the letter grade scheme, the quality of evidence for these trials is C+.

**Figure 1.**
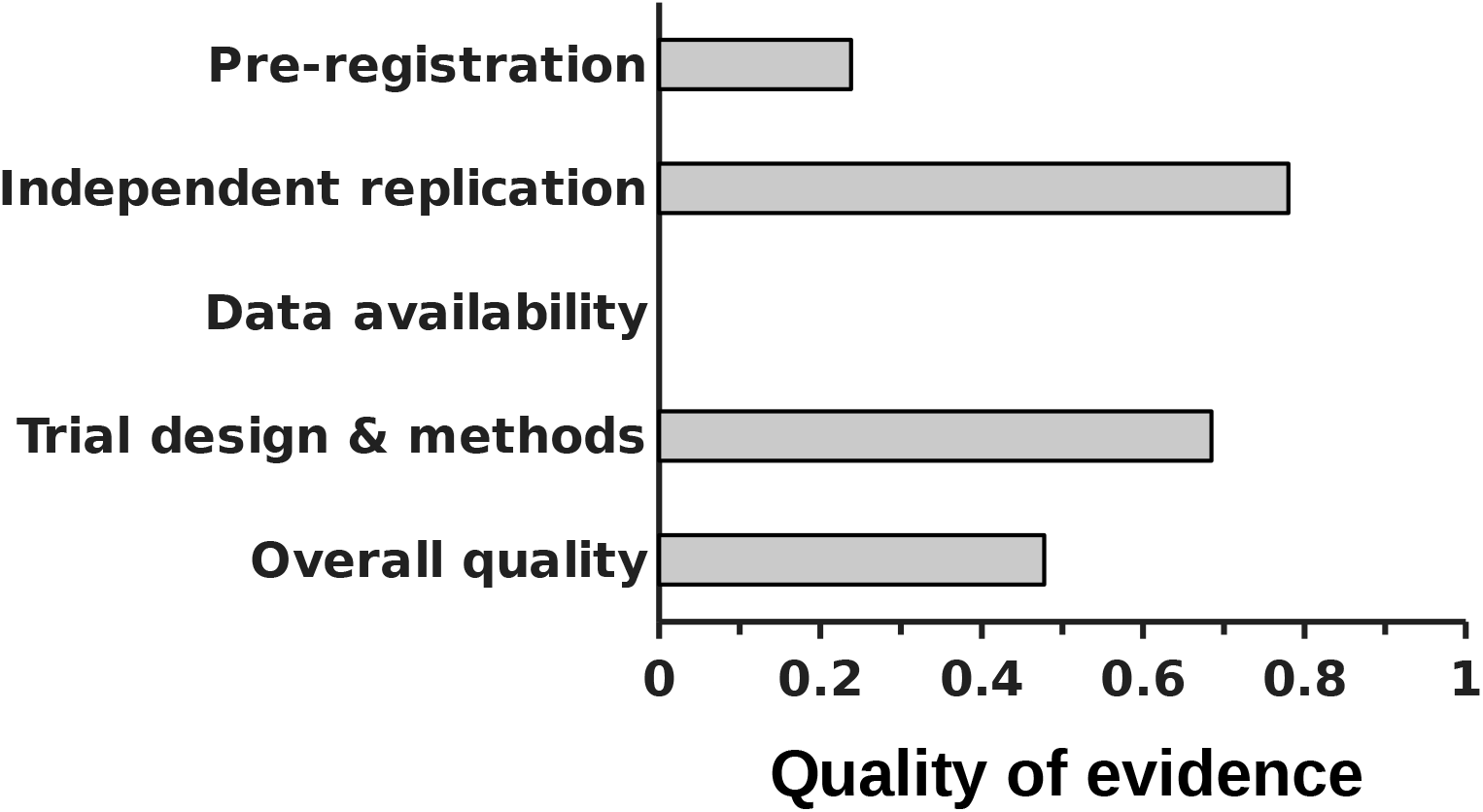
Quality of evidence for specific dimensions and overall for trials summarized by Machado and colleagues (14)

## Discussion

The index of quality enables systematic, quantitative measurement of this key facet of evidence in meta-analysis of RCTs. As such, it is a counterpart to other fundamental summary measures in meta-analysis. The sub-indexes of pre-registration, independent replication, data availability, and trial design and methods can also be used separately or in combination with the overall quality index to describe the multi-dimensional nature of the quality of evidence. Several of these sub-indexes could also be applied to meta-analyses of observational studies. For illustration, I computed the quality index for the RCTs summarized in a recent meta-analysis, whose authors rated the evidence as “high quality” based on the unsystematic GRADE procedures that do not cover most of the dimensions in the quality index. These trials have only a moderate (C+) level of quality when assessed systematically with the quality index.

Analysts may vary in their coding of the inputs to the quality index. Researchers concerned about this potential source of error can employ multiple raters and estimate their inter-rater reliability. Better yet, those applying this index can simply report the data underlying their calculation of the index. Other researchers could then verify, challenge, or recalculate index values for the same trials.

The standards for conducting and reporting randomized controlled trials vary across fields and have changed over time. The quality scores, therefore, are conservative–possible underestimates–especially concerning interventions for which much of the evidence is based on older (say, before 2000) trials, because authors may have actually met some of the quality standards, but just did not report enough details to confirm that they did. Also, some practices, such as pre-registration, were almost non-existent until recently. Older norms and practices may have impacts on the quality of evidence for many topics for some time, at least until newer evidence gathered with higher standards accumulates substantially.

Some reviewers contend that absence of reporting a trial design and methods feature should not be mistaken as absence of that feature in assessing quality (2, 21). While it is impossible to know whether lack of reporting represents absence of the feature, reviewers can synthesize only the evidence that has been reported. There is no universally applicable alternative approach for summarizing the evidence objectively. In some cases, meta-analysts can try to contact authors to get more completely information. Although this is desirable, it is often not possible to do (for older studies) or fails to yield the necessary information.

Quality of evidence may vary unavoidably among types of interventions. For example, trials of interventions for which blinding is impossible may tend to have lower trial design quality than for interventions that can be evaluated with blinding.

With the exception of one part of the pre-registration component, the quality of evidence index is based entirely on information about the trials included in a meta-analytic summary. Although there may be no information external to the trials that can be used to validate them, there may be at least one circumstance in which external information could cast doubt on the evidence from the trials. When RCTs show a strongly positive intervention effect on average, but observational studies (such as cohort, case-control, and cross-sectional studies) show a negative or negligibly positive association with the intervention, the results of the RCTs may sometimes be considered suspect. Observational evidence is typically the warrant for conducting an RCT. If observational studies tend to indicate negative or negligibly positive results, suggest no obvious confounders, and precede the RCTs, then the integrity of RCT evidence showing strongly positive results may be questionable.

The quality of evidence index does not include several factors that some researchers might consider aspects of quality. The research funding source does not indicate how the evidence was actually produced. Yet, if funding source were reliably associated with estimated effects—a true bias—it might be reasonable to consider it an aspect of quality. Many observers are concerned about industry sponsorship specifically, hypothesizing that such funding leads researchers to generate results that favor the sponsor’s product or service. However, there seems to be no consistent relationship between industry sponsorship and effect size in RCTs. In November, 2016, I examined reports cited in Lundh and colleagues’ systematic review of the topic (22) and sources citing their review, as indicated by Google Scholar (focusing on those with relevant titles; scholar.google.com). I found four meta-analyses that assessed the association between industry sponsorship and effect size for trials in people with the same intervention, comparator, indication, and outcome. Associations ranged from slightly negative to mildly positive (23–26). Nonetheless, funding source is still a relevant dimension to summarize separately from quality, as it may affect trust in results.

Generalizability of results and overall attrition also are not factors in the quality index. Quality is a characteristic *internal* to completed studies. Generalizability is a judgment about how well participants in completed studies correspond to a particular population *external* to the completed studies (2). Moreover, this population and its defining characteristics may vary across researchers and time. Overall attrition relates to generalizability. If attrition in an RCT is high, but similar between arms, then participants completing the study might not representative of the whole initial sample and the estimated effect could differ from one that might have been observed had attrition been low. Participant characteristics, trial duration, study procedures, and other factors could affect overall attrition. These factors also influence recruitment to RCTs. Thus, overall attrition may produce genuine variability in the estimated effect between trials in the same way that differences between trials in pre-randomization participant characteristics do.

Each of the quality of evidence sub-indexes has limitations. For the pre-registration measure, the timing of the review and delayed reporting may affect the proportion of pre-registered, completed trials that have not been reported. That is, some old trials may have been reported, but after more than three years since completion. These old trials would be included in the review, but might not have been included had the review been conducted earlier. In this situation, the proportion of pre-registered trials that had been reported would be lower at the earlier time point.

The independent replication sub-index could be misestimated when multiple individuals with the same full name publish different reports that are summarized in the same meta-analysis. Several author identification services exist currently but none cover a significant proportion of authors well. Even if a service were to become standard and comprehensive in the future, it would not cover past authors who no longer publish. Also, my definition of independent teams of authors–none in common–has face validity and enables objective assessment. However, it could under- or over-state independence in some cases. Furthermore, this sub-index does not capture any tendency toward conformity among researchers, even for those working independently from each other. Ioannidis (27) hypothesized that authors may produce results to align with prevailing beliefs and that editors and reviewers may shape the literature to reinforce dominant views. Nationalization of research funding, with peer review as the key evaluative component, may fuel some of this groupthink (28).

The data availability sub-index is a passive indicator because it relies entirely on authors explicitly noting the availability of their data in a public archive or publishing them in their report. Currently there are many scientific data archives and no centralized database for searching most or all of them as there is for clinical trial registrations (the International Clinical Trials Registry Platform). Analysts could choose, however, to designate particular data archives they will search for supplementing the information about data availability authors give in their reports. This measure also does not capture the negative end of the data availability spectrum: when authors refuse to share accessible data even when specifically asked (29, 30). If authors refuse to share data in such circumstances, the study may be considered especially suspect and not worthy of inclusion in a systematic review or meta-analysis.

The trial design and methods sub-index involves passive factors as well. For instance, early stopping and the use of randomization units other than individual participants, in particular, may not be detectable unless authors explicitly note them. Authors seldom specify that participants were *individually* randomized (except when mentioning some sort of blocking or related procedure), so meta-analysts generally must assume randomization occurred at the individual level. My arbitrary threshold for differential attrition (absolute 10% difference) is dichotomous and hence does not distinguish between moderate and large degrees of differential attrition. Similarly, this factor does not account for differential time to dropout (13). I excluded this as a factor because authors rarely report it and it is likely strongly related to (and thus redundant with) the differential dropout measure. Finally, in this sub-index, each trial design and methods factor has an arbitrary equal weight.

Eventually, for new research topics, it may be possible to restrict systematic reviews and meta-analyses just to pre-registered RCTs. For reviews that involve “legacy” literature, though, pre-registration will remain an important quality criterion. Ideally, all phases of the research process–not just the protocol at the beginning and publicly archived data at the end–would be transparent, auditable, and tamper-proof. The technology to make this easy to do is not yet available. If this ability materializes in the future, the quality of evidence index could be updated to incorporate the use of such tools as an additional dimension. Still, science might forever be based on the “honor system,” as ultimately researchers have to trust, to a large extent, the findings of their peers. Unfortunately, there might always be a way for unscrupulous or careless researchers to subvert systems that promote transparency. Therefore, perpetual vigilance and preoccupation with research quality may be required to reduce error in the scientific record.

## Acknowledgments

I thank Britton Brewer, Stuart Brody, Barbara Leigh, John Potterat, and John Roberts, Jr. for their helpful comments on an earlier draft of this paper. All remaining errors and omissions are my own. I received no funding for this work and I declare no conflicts of interest.

